# Spike protein cleavage-activation in the context of the SARS-CoV-2 P681R mutation: an analysis from its first appearance in lineage A.23.1 identified in Uganda

**DOI:** 10.1101/2021.06.30.450632

**Authors:** Bailey Lubinski, Laura E. Frazier, My V.T. Phan, Daniel L. Bugembe, Jessie L. Cunningham, Tiffany Tang, Susan Daniel, Matthew Cotten, Javier A. Jaimes, Gary R. Whittaker

## Abstract

Based on its predicted ability to affect transmissibility and pathogenesis, surveillance studies have highlighted the role of a specific mutation (P681R) in the S1/S2 furin cleavage site of the SARS-CoV-2 spike protein. Here we analyzed A.23.1, first identified in Uganda, as a P681R-containing virus several months prior to the emergence of B.1.617.2 (Delta variant). We performed assays using peptides mimicking the S1/S2 from A.23.1 and B.1.617 and observed significantly increased cleavability with furin compared to both an original B lineage (Wuhan-Hu1) and B.1.1.7 (Alpha variant). We also performed cell-cell fusion and functional infectivity assays using pseudotyped particles and observed an increase in activity for A.23.1 compared to an original B lineage spike. However, these changes in activity were not reproduced in the B lineage spike bearing only the P681R substitution. Our findings suggest that while A.23.1 has increased furin-mediated cleavage linked to the P681R substitution, this substitution needs to occur on the background of other spike protein changes to enable its functional consequences.

## Introduction

Severe acute respiratory syndrome coronavirus 2 (SARS-CoV-2) is the agent causing the current COVID-19 pandemic (1). SARS-CoV-2 was first identified in late 2019 and has since spread rapidly throughout the world. The virus emerged as two main lineages, A and B, and now multiple sub-lineages. While the B.1 lineage became the dominant virus following its introduction into Northern Italy and spread through Europe/UK in February 2020, both A and B lineages remain in circulation globally (2). Both lineages have undergone significant diversification as they expanded; this expansion is apparently linked to a key S gene mutation—D614G in lineage B.1 and all sublineages, which has been linked to modest increase in virus transmissibility (3) and with Q613H found in lineage A.23/A.23.1. As Q613H is adjacent to D614G it may represent an example of convergent evolution that resulted in a more stabilized spike protein (4). D614G has now become established in circulating B and derived lineages. Compared with the lineage B.1 viruses that have successfully evolved into multiple variants of concern (VOC), including B.1.1.7 (Alpha) B.1.351 (Beta), B.1.1.28.1/P.1 (Gamma), B.1.617.2 (Delta), most of lineage A viruses remained at fairly lower frequency and were more prevalent at the beginning of the pandemic in Asia. However, the A.23.1 viral lineage is one of a few lineage A viruses that, due to local circumstances, became abundant in Uganda (5), Rwanda (6) and South Sudan (7). A.23.1 evolved from the A.23 virus variant first identified in Uganda in July 2020 and is characterized by three spike mutations F157L, V367F and Q613H (5). Subsequently, the evolving A.23.1 lineage acquired additional spike substitutions (P681R), as well as in nsp6, ORF8 and ORF9 and with the acquisition of the E484K substitution A.23.1 was designated a variant under investigation (VUI). By July 2021, the A.23.1 lineage has been observed with 1110 genomes reported from 47 countries (GISAID, Pango Lineage report; https://cov-lineages.org/global_report_A.23.1.html).

Among several mutations in the A.23.1 lineage, the P681R mutation is of interest as it is part of a proteolytic cleavage site for furin and furin-like proteases at the junction of the spike protein receptor-binding (S1) and fusion (S2) domains (8). The S1/S2 junction of the SARS-CoV-2 S gene has a distinct indel compared to all other known SARS-like viruses (Sarbecoviruses in *Betacoronavirus* lineage B)—the amino acid sequence of SARS-CoV-2 S protein is _681_-P-R-R-A-R|S-_686_ with proteolytic cleavage (|) predicted to occur between the arginine and serine residues depicted. Based on nomenclature established for proteolytic events (9), the R|S residues are defined as the P1|P1’ residues for enzymatic cleavage, with residue 681 of A.23.1 spike being the P5 cleavage position. The ubiquitously-expressed cellular serine protease furin is highly specific and cleaves at a distinct multi-basic motif containing paired arginine residues; furin requires a minimal motif of R-x-x-R (P4-x-x-P1), with a preference for an additional basic (B) residue at P2; i.e., R-x-B-R (10). For SARS-CoV-2, the presence of the S1/S2 “furin site” enhances virus transmissibility (11, 12). For the A.23.1 S, P681R provides an additional basic residue at P5 and may modulate S1/S2 cleavability by furin, and hence virus infection properties (13). Notably, the P681R substitution appears in several other lineages, most notably B.1.617.2 (Delta) (N = 617590 genomes) but also the AY.X sub-lineages of B.1.617.2, B.1.617.1 (N= 6138), B.1.466.2 (N=2208), the B.1.1.7 sub-lineage Q.4 (2067), B.1.551 (N=722), AU.2 (N=302), B.1.1.25 (N=509), B.1.466.2 (N=538), and other lineages (updated 22 Oct 21, https://outbreak.info; https://outbreak.info/situation-reports?pango&muts=S%3AP681R) suggesting that the substitution may provide an advantage for viruses encoding the substitution.

We previously studied the role of proteolytic activation of the spike protein of the lineage B SARS-CoV-2 isolates Wuhan-Hu1 and B.1.1.7 (14). Here, we used a similar approach to study the role of the proteolytic activation of the spike protein in the context of the A.23.1 lineage virus, with a focus on the P681R substitution to better understand the role of this notable change at the S1/S2 (furin) cleavage site.

## Results

### Emergence and analysis of SARS-CoV-2 variants in Uganda, and evolution of the P681R mutation and its role in the transmissibility and emergence of SARS-CoV-2

A summary of the daily reported SARS-CoV-2 infections in Uganda is shown in Figure 1A, along with a summary of SARS-CoV-2 lineage data in samples from Uganda (Figure 1B). The peak of infections in December 2020-January 2021 corresponded to the circulation of the A.23.1 variant, which subsided, but was replaced by a second larger peak of infections beginning in July 2021, primarily due to the emergence of the B.1.617.2 variant (Delta), with additional variants being present over time. The circulation of the A.23.1 variant appears to be fully displaced by the B.1.617.2 variant, which by July 2021 became the prevalent variant in this country.

**Figure 1:**
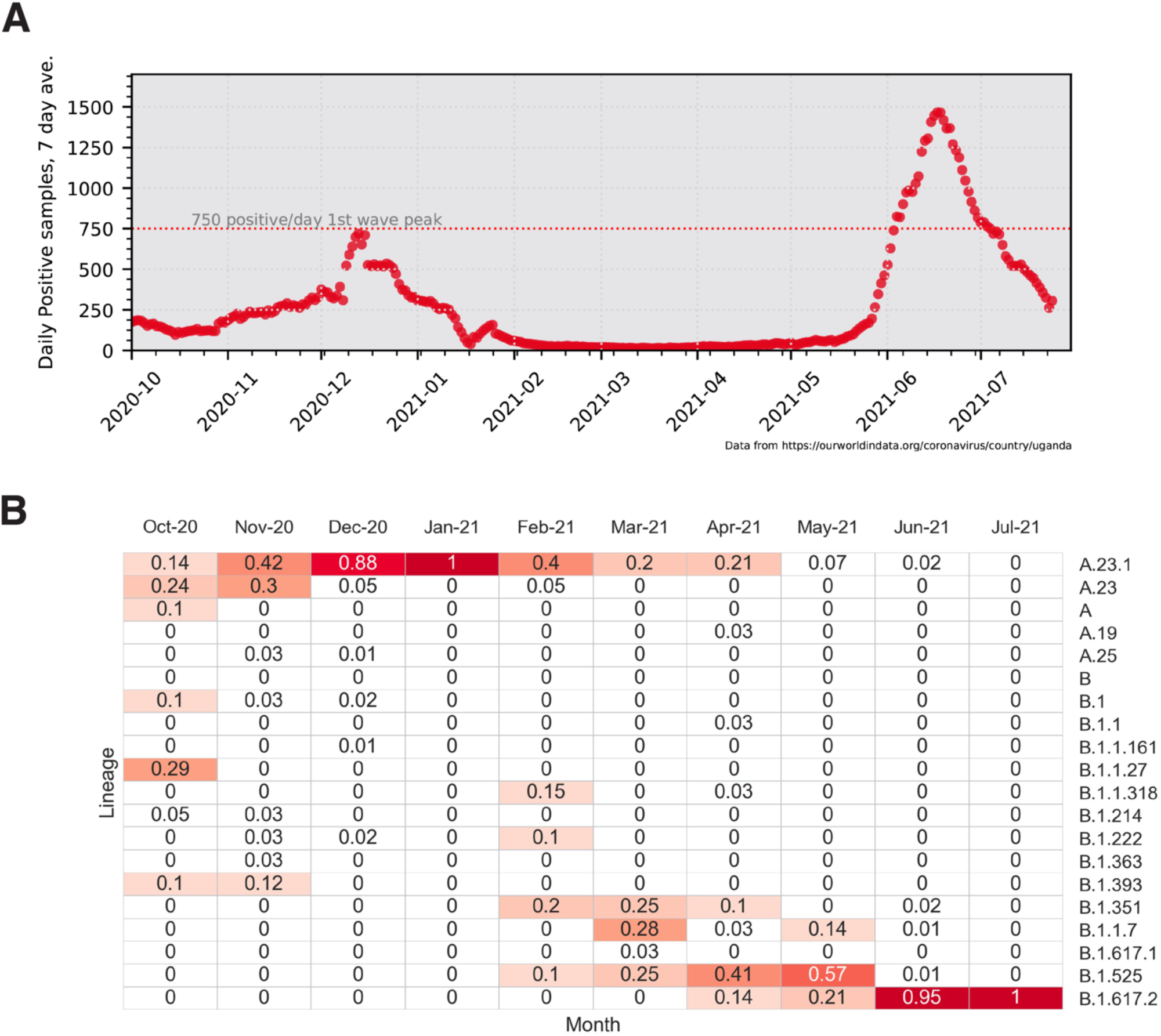
Uganda SARS-CoV-2 cases and lineages, October 2020 to July 2021. **A**. 7-day average positive cases numbers were plotted by day, the peak of 750 cases/ per day observed in the first wave of infections in January 2021 is indicated with a dotted line. Case data were obtained from Our World in Data (https://ourworldindata.org/). **B**. Monthly SARS-CoV-2 lineage data for Uganda. All Uganda full genome sequences from GISAID (https://www.gisaid.org/) were retrieved, lineage types using the Pango tool (https://cov-lineages.org/resources/pangolin.html), and the fraction of each month’s total genomes were plotted. Fractions were indicated in each cell and cells are colored (white to dark red) by increasing fraction.

To further understand the evolution of the P681R substitution and its role in the transmissibility of SARS-CoV-2, we monitored the frequency of substitutions at the S1/S2 (furin cleavage) site in the global surveillance data and plotted these substitution data as fraction of total genomes. The initial B lineage virus that spread out of Wuhan encoded a spike protein with P681 at the furin cleavage site, along with G614. Fairly early in the epidemic, the D614G substitution appeared in the B.1 lineage and became prevalent in May-December 2020 (Figure 2A). The B.1.1.7 (Alpha) lineage evolved from B.1, spread widely in the UK and other regions, and encoded a P681H substitution in the G614D background. B.1.1.7 peaked globally in March-April 2021 (Figure 2A). In most regions of the world, the B.1.617.2 (Delta) lineage encoding spike D614G and P681R spread widely following its emergence in India in May-June 2021, and became the dominant observed lineage globally (Figure 5A). In comparison, a distinct A lineage virus (A.23) containing Q613H appeared in August-September 2020 and acquired the P681R mutation at a much earlier time (December 2020-January 2021); however, it circulated only briefly (Figure 2B). These data suggest that while P681R is important, it operates in the context of other viral mutations in the context of community spread, with this functional context able to be addressed experimentally.

**Figure 2.**
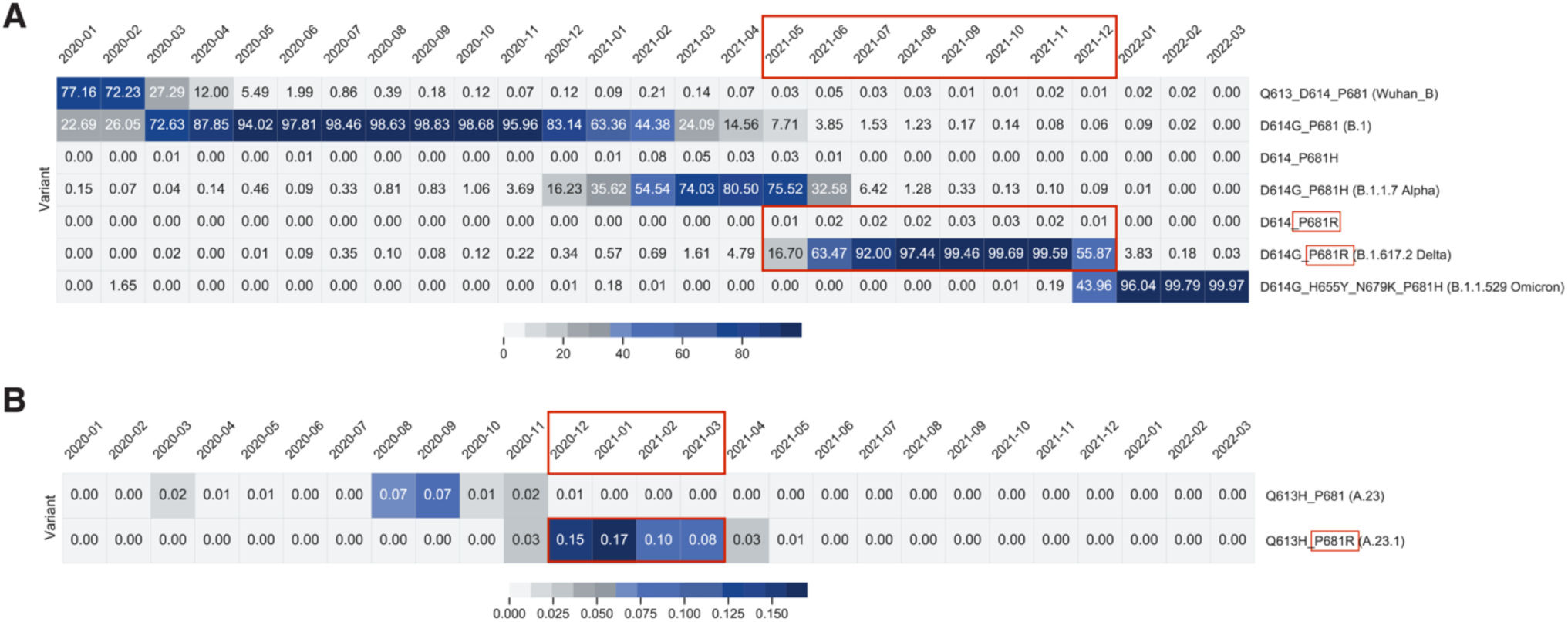
Frequency of P681, P681R, P681H, D614 D614G and D613H substitutions. The frequency of substitutions was counted by string matching to a peptide sequencing spanning the position 613 to 681) including the relevant sites at 613/614 and 681. (D614_P681 (Wuhan_B), D614G_P681 (B.1), D614_P681H, D614G_P681H (B.1.1.7), D614_P681R, Q613_P681 (Wuhan_B), Q613H_P681 (A.23), Q613H_P681R (A.23.1), D614G_P681R (B.1.617.2). Fractions of total genomes available for each month were plotted. Color bar at the bottom of each panel indicate fraction/color code. **A**. Lineage B relevant substitutions. **B**. Lineage A.23 and A.23.1 relevant substitutions. The time periods where P681R was dominant in each lineage are shown in red boxes.

### Biochemical analysis of the SARS-CoV-2 A.23.1 S1/S2 cleavage site

To gain insight into SARS-CoV-2 spike protein function and the proteolytic processing at the S1/S2 site, we took a combined biochemical and cell-based strategy, with the rationale that along with other changes in the spike protein, A.23.1, B.1.617.1 (Kappa) and B.1.617.2 (Delta) contain a P681R substitution at the S1/S2 interface which may modulate spike protein function and that these mutations alter the furin cleavage site—which can be monitored by analyzing downstream changes in the levels of cleaved products and in virus-cell fusion and pseudoparticle activation. Sequences of representative S1/S2 sequences are summarized in Figure 3A.

**Figure 3:**
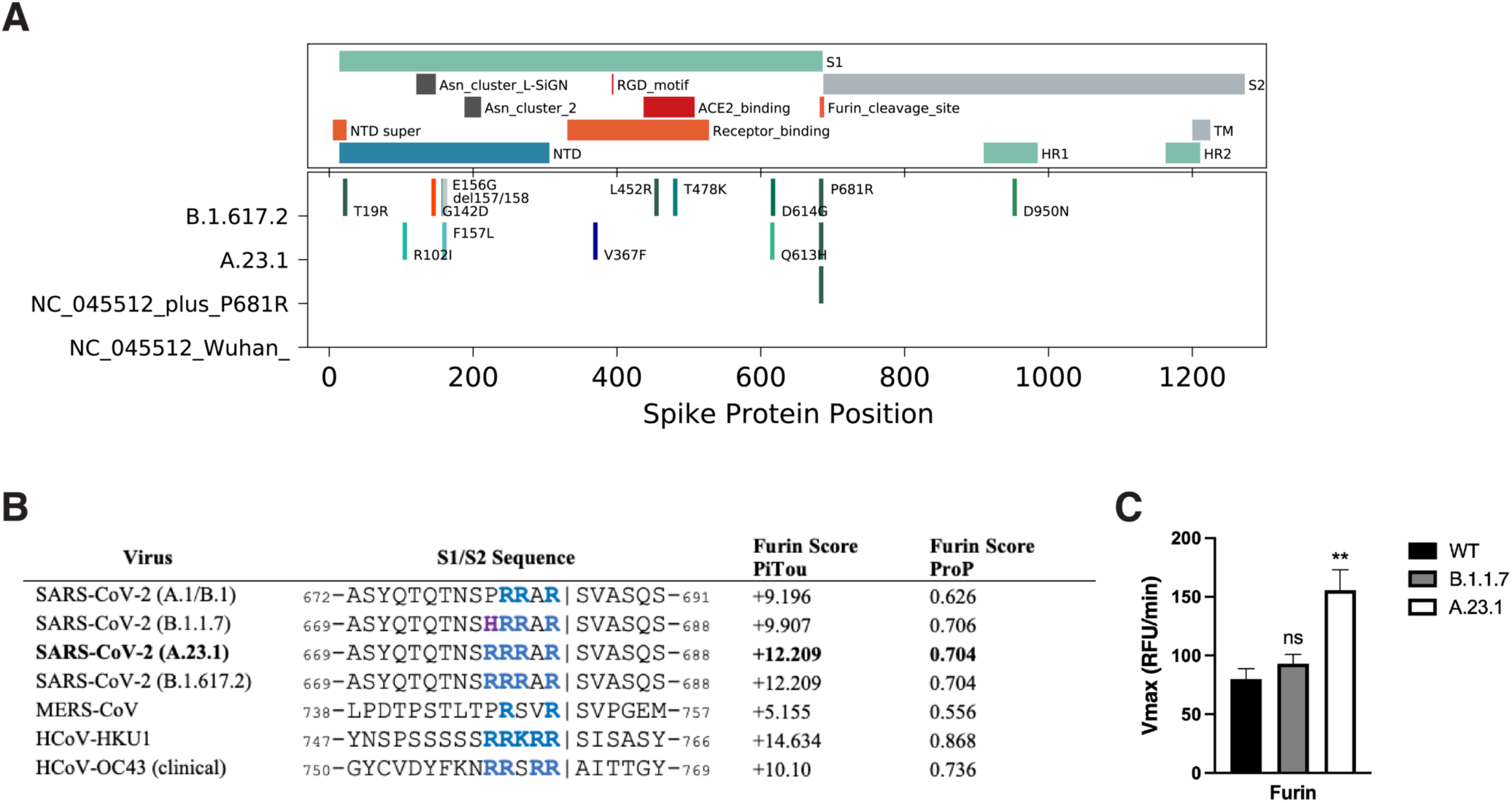
SARS-CoV-2 A.23.1 S sequence changes and S1/S2 furin cleavage. **A**. Summary of notable functional domains and sequence changes in the spike gene of A.23.1 compared to Wuhan-Hu-1 and B.1.617.2 (Delta). **B**. Furin cleavage score analysis of CoV S1/S2 cleavage sites. CoV S sequences were analyzed using the ProP1 1.0 and PiTou2 3.0 furin prediction algorithm, generating a score with bold numbers indicating predicted furin cleavage. (|) denotes the position of the predicted S1/S2 cleavage site. Basic resides, arginine (R) and lysine (K), are highlighted in blue, with histidine in purple. Sequences corresponding to the S1/S2 region of SARS-CoV-2 (QHD43416.1), SARS-CoV (AAT74874.1), MERS-CoV (AFS88936.1), HCoV-HKU1 (AAT98580.1), HCoV-OC43 (KY369907.1) were obtained from GenBank. Sequences corresponding to the S1/S2 region of SARS-CoV-2 B.1.1.7 (EPI_ISL_1374509) and SARS-CoV-2 A.23.1 hCoV-19/Uganda/UG185/2020 (EPI_ISL_955136), were obtained from GISAID. **C**. Fluorogenic peptide cleavage assays of the SARS-CoV-2 S1/S2 cleavage site. Peptides mimicking the S1/S2 site of the SARS-CoV-2 Wuhan-Hu-1 (WT – P681), B.1.1.7 (P681H) and A.23.1 (P681R) variants were evaluated for in vitro cleavage by furin, compared to trypsin control. Error bars represent G standard errors (n = 9). Asterisks indicate statistical significance compared to the untreated control. Statistical analysis was performed using an unpaired Student’s t test. ** p < 0.01.

As an initial bioinformatic approach to assess biochemical function, we utilized the PiTou (15) and ProP (16) protein cleavage prediction tools, comparing the spike proteins from A.23.1 to B.1.1.7 and the prototype SARS-CoV-2 from the A.1 and B.1 lineages (e.g., Wuhan-Hu-1), as well as to MERS-CoV and selected other human respiratory betacoronaviruses (HCoV-HKU1 and HCoV-OC43) with identifiable furin cleavage sites (Figure 3B). Both algorithms predicted an increase in the furin cleavage for the A.23.1 and B.1.617 lineages compared to A.1/B.1, with PiTou also showing a marked increase compared to B.1.1.7. PiTou utilizes a hidden Markov model specifically targeting 20 amino acid residues surrounding furin cleavage sites and is expected to be a more accurate prediction tool. As expected, MERS-CoV showed a relatively low furin cleavage score, with HCoV-HKU1 and HCoV-OC43 showing relatively high furin cleavage scores. Overall, these analyses predict a distinct increase of furin cleavability for the spike protein of A.23.1 and B.1.617 lineage viruses compared to A.1/B.1 and B.1.1.7. lineage viruses.

To directly examine the activity of furin on the SARS-CoV-2 A.23.1 S1/S2 site, we used a biochemical peptide cleavage assay to directly measure furin cleavage activity *in vitro* (17). The specific peptide sequences used here were SARS-CoV-2 S1/S2 B.1.1.7 (TNSHRRARSVA), TNSPRRARSVA (Wuhan-Hu-1 S1/S2) and TNSRRRARSVA (A.23.1 S1/S2). As predicted, furin effectively cleaved both the Wuhan-Hu-1 (WT) and B.1.1.7 peptides, but with no significant differences (Figure 3C). Interestingly, and agreeing with the PiTou prediction, we observed a significant increase in furin cleavage for the A.23.1 S1/S2 peptide (Figures 2B and C) compared to both Wuhan-Hu-1 (WT) and B.1.1.7. This comparative data with SARS-CoV S1/S2 sites reveals that the P681R substitution substantially increases cleavability by furin, beyond the small increase noted previously for P681H (11).

### Cell-to-cell fusion assays of A.23.1 spike

In order to assess the functional properties of the spike protein and to see if the P681R substitution provided any advantage for cell-to-cell transmission or syncytia formation, we performed a cell-to-cell fusion assay in which VeroE6 or Vero-TMPRSS2 cells were transfected with either the WT, A.23.1 or P681R spike gene. We then evaluated syncytia formation as a read-out of membrane fusion. We observed an increase in the syncytia formation following spike protein expression for either A.23.1 or Wuhan-Hu-1 harboring a P681R mutation (P681R), compared to Wuhan-Hu-1 (WT) (Figure 4A). Vero-TMPRSS2 cells generally formed more extensive syncytia than VeroE6 cells. This increase was evident by observation through fluorescence microscopy, as well as by quantification of the syncytia and cell-to-cell fusion ratio (Figure 4A, B and C). An increase in the number of nuclei involved in syncytia was observed in cells transfected with A.23.1 and P681R S genes in both cell lines (Figure 4B). However, the increase was higher in Vero-TMPRSS2 cells in all the three studied spike proteins. Membrane expressed spike cleavage also assessed using western blot (Figure 4C). An increased cleavage ratio was observed in the A.23.1 and P681R membrane expressed spikes, compared to WT. The cleavage ratio was similar in both cell lines. Band intensity was normalized to the GLUT4 protein (housekeeping protein) band intensity. These data provide evidence that the P681R mutation increases membrane fusion activity of the SARS-CoV-2 spike protein under the conditions tested.

**Figure 4:**
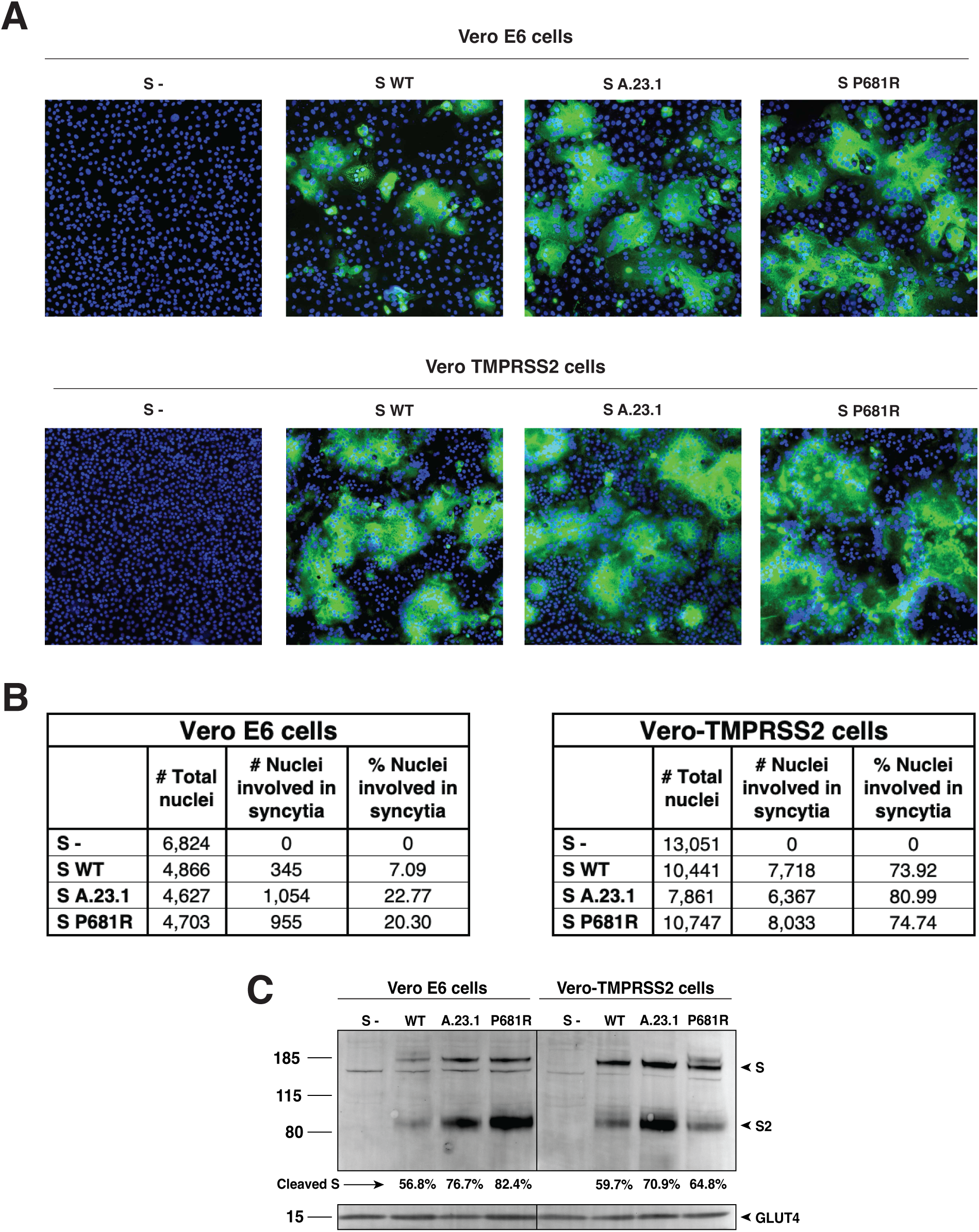
Cell-to-cell fusion in SARS-CoV-2 A.23.1 S expressing cells. **A**. Cell-to-cell fusion assay of SARS-CoV-2 Wuhan-Hu-1 S (WT), SARS-CoV-2 S A.23.1 variant, or SARS-CoV-2 S WT with P681R mutation. S- = non-transfected cells. SARS-CoV-2 S was detected using a rabbit antibody against the SARS-CoV-2 S2 region. **B**. Syncytia quantification by number of nuclei involved in syncytia. **C**. Western blot analysis of membrane expressed S proteins and GLUT4 (housekeeping expression protein). All the experiments were performed on Vero E6 and Vero-TMPRSS2 cells.

### Functional analysis of virus entry using viral pseudoparticles

To assess the functional importance of the S1/S2 site for SARS-CoV-2 entry, we utilized viral pseudoparticles consisting of a murine leukemia virus (MLV) core displaying a heterologous viral envelope protein to partially recapitulate the entry process of the native coronavirus. The pseudoparticles also contain a luciferase reporter gene as well as the integrase activity to allow that integration into the host cell genome to drive expression of quantifiable luciferase (MLVpp-SARS-CoV-2 S) (18). Using the HEK-293T cell line for particle production, MLV pseudoparticles containing the spike proteins of A.23.1, Wuhan-Hu-1 SARS-CoV-2 (WT), and a P681R point mutant of Wuhan-Hu-1 (P681R) were prepared. Positive-control particles containing the vesicular stomatitis virus (VSV) G protein and negative-control particles (Δenvpp) lacking envelope proteins were also prepared (not shown). Pseudoparticles were probed for their S content via western blot (Figure 4A). Because SARS-CoV-2 S has an efficiently cleaved spike we also produced particles under furin inhibition using dec-RVKR-CMK. This allowed changes in cleavage patterns between different spike proteins to be visualized. We also treated the particles with exogenous furin (+ or – furin) to examine the spike cleavage on both partially cleaved and uncleaved spikes, to further study the differences in furin processing in the studied spikes. For both A.23.1 and P681R particles, we detected increased spike protein cleavage compared to WT in the harvested particles (Figure 5A). Interestingly, we observed markedly increased cleavage ratio for both A.23.1 and P681R spikes in the harvested pseudoparticles under furin-inhibition conditions (Figure 5A), presumably as the enhanced furin cleavage motif site produced by the P681R mutation rescued what may be a modest effect of dec-RVKR-CMK.

**Figure 5:**
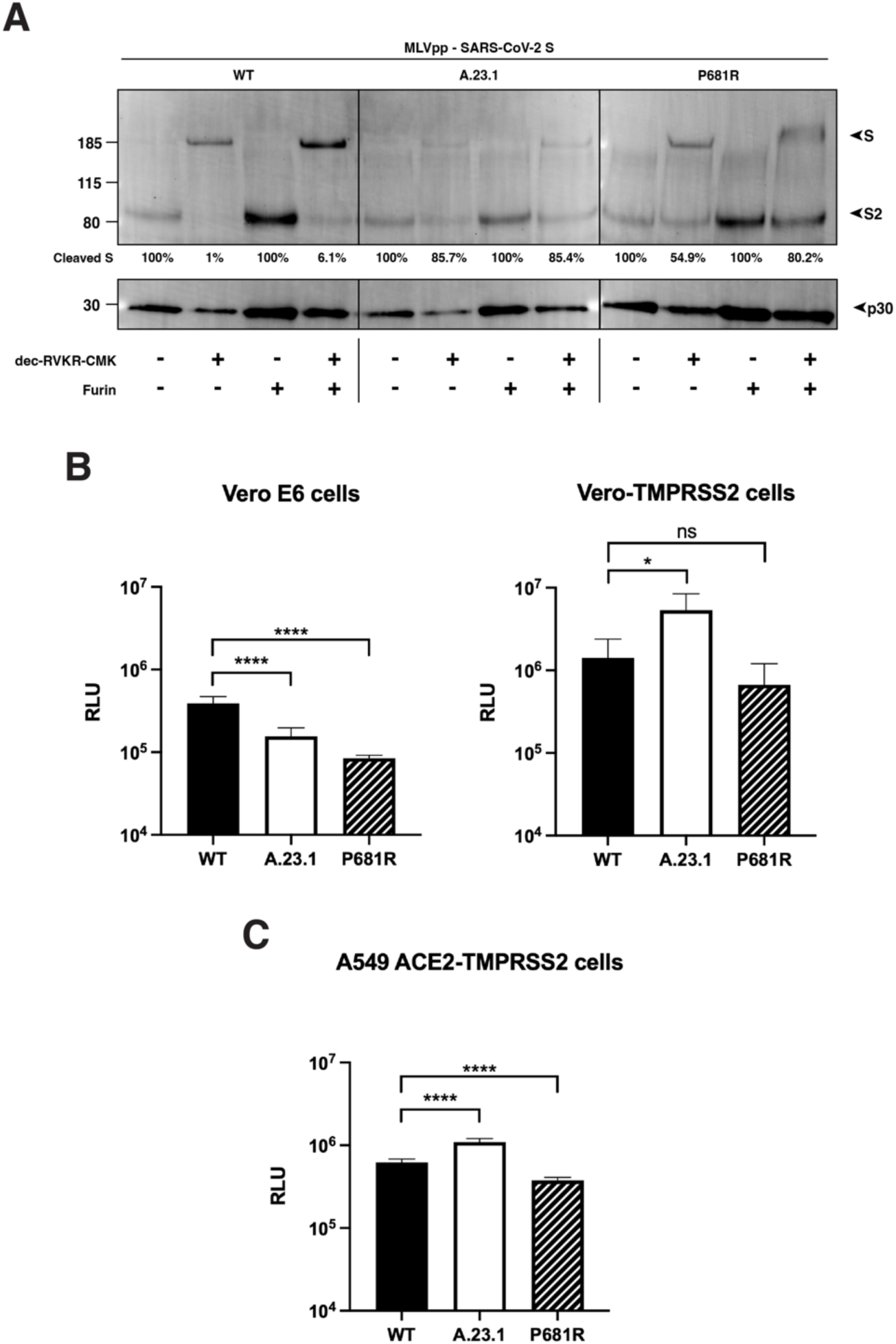
SARS-CoV-2 A.23.1 variant S1/S2 cleavage site activation and role in viral entry. **A**. Western blot analysis of MLVpp-SARS-CoV-2 S produced in ± dec-RVKR-CMK and treated with ± furin. S was detected using a rabbit antibody against the SARS-CoV-2 S2 subunit. MLV content was detected using a mouse antibody against MLV p30. **B**. Pseudoparticle infectivity assays in Vero E6 and Vero-TMPRSS2 cells. Cells were infected with MLVpps harboring the VSV-G, SARS-CoV-2 S (WT), SARS-CoV-2 S A.23.1 variant, SARS-CoV-2 S WT with P681R mutation. Data represents the average luciferase activity of cells of four independent experiments (Vero E6 and Vero-TMPRSS2). Error bars represent G standard deviation (n = 3). Asterisks indicate statistical significance compared to the untreated control. Statistical analysis was performed using an unpaired Student’s t test. * p < 0.1, **** p < 0.0001. **C**. Pseudoparticle infectivity assays in A549-ACE2-TMPRSS2 cells. Cells were infected with MLVpps harboring the VSV-G, SARS-CoV-2 S (WT), SARS-CoV-2 S A.23.1 variant, SARS-CoV-2 S WT with P681R mutation. Data represents the average luciferase activity of cells of three independent experiments. Error bars represent G standard deviation (n = 3). Asterisks indicate statistical significance compared to the untreated control. Statistical analysis was performed using an unpaired Student’s t test. **** p < 0.0001.

For SARS-CoV-2, furin is predicted to cleave during virus assembly and “prime” the spike protein at the S1/S2 site for subsequent events during cell entry. However, a subsequent cleavage priming at a secondary site (known as S2’ site) is also needed to activate the spike’s fusion machinery (1). SARS-CoV-2 is predicted to enter Vero E6 cells using cathepsin L for activation during endosomal trafficking, in what is known as the “late” pathway, whereas in Vero-TMPRSS2 is predicted to use a “early” pathway, with spike activated by TMPRSS2 or other transmembrane serine proteases (TTSPs) at the cellular membrane (1). Here, we used the Vero-TMPRSS2 and the Vero E6 cell lines, which are predicted to activate the SARS-CoV-2 using TMPRSS2 and cathepsin L respectively. Considering that furin priming at the S1/S2 site normally occurs during viral assembly, we used pseudoparticles that were produced without furin inhibitor, yielding cleaved spike proteins. Vero-TMPRSS2 cells gave overall significantly higher luciferase signals indicative of more efficient entry. In contrast, Vero E6 cells showed generally lowered infection levels. As expected, VSVpp (positive control) pseudoparticles infected both cell lines with several orders of magnitude higher luciferase units than the values reported with Δenvpp infection (data not shown). In Vero E6 cells, entry of A.23.1. and P681R was lowered compared to wild type (Figure 5B). However, Vero-TMPRSS2 cells pseudoparticles bearing the A.23.1 spike showed a significantly higher level of infection, indicating more efficient virus entry; This was not reproduced for a P681R point mutant of Wuhan-Hu-1 (P681R), a result in line with previous results indicating that other mutations in spike are needed for the increased cleavability imparted by the P681R mutation to mediate enhanced virus infection.

As a further way to assess the entry mediated by A.23.1. and P681R-containing spike proteins, we tested pseudoparticles in human lung A459 cells expressing ACE2 and TMPRSS2 (Figure 5C). These cells showed a highly significant increase in infection by A.23.1 compared to Wuhan-Hu1. The point mutant of P681R on the Wuhan-Hu1 background showed a decrease in infectivity, confirming that P681R (similarly to other point mutants (19)) only has its functional consequence on the appropriate genetic background.

## Discussion

Since late 2020, the evolution of the SARS-CoV-2 pandemic has been characterized by the emergence of viruses bearing sets of substitutions/deletions, designated “variants of concern” (VOCs) and “variants under investigation” (VUIs). These variants appear to have expanded following the selection for substitution or deletions in the spike protein, such as D614G and Q613H, along with mutations in other viral proteins. The substitutions encoded by such variants may alter virus characteristics including enhanced transmissibility and antigenicity, some provide a direct advantage to avoid the changing developing immune responses in the population due to prior exposure or vaccination as well as the social dynamics of the human population (4, 20-23). The specific case of the D614G is interesting, as this mutation have been shown to improve the spike’s open conformation for receptor binding, demonstrating an evolutionary advantage for the 614G carrier virus (24). In fact, the explosive spread of COVID-19 cases can be tracked to the emergence of this mutation, which provided the context for further evolution of the SARS-CoV-2 virus and the rising number of new variants. The first notable SARS-CoV-2 VOC of 2021 was B.1.1.7 (Alpha), which among other changes, encoded a P681H substitution in the spike S1/S2 furin cleavage site and has been linked to increased transmissibility due to the presence of the additional basic amino acid, histidine (H). However, histidine is unusual in that it has an ionizable side chain with a p*K*a near neutrality (25), and so is not conventionally considered a basic amino acid. Most recently, the VOC (B.1.617.2, or Delta) has replaced B.1.1.7(Alpha) as the dominant circulating virus globally, which like A.23.1 and sub-lineages B.1.617.1, B.1.617.2 and B.1.617.3 encodes a P681R substitution, and is more conventionally “polybasic” in the S1/S2 cleavage motif, than the P681H of B.1.1.7 (Alpha) and is suggested to affect transmissibility and pathogenesis (26). For the Delta variant (B.1.617.2), Saito *et al*. showed enhanced fusogenicity and viral entry in cells expressing TMPRSS2 (Vero-TMPRSS2 and Calu-3) but lowered fusogenicity in Vero cells (27), with equivalent results also shown by Peacock et al. in a range of TMPRSS2-expressing cells (28). Our data with Vero cells in particular differ from those reported by Saito *et al*., reinforcing the concept that cell-cell fusion can be affected by many factors, including the specific growth conditions of the cells, and also suggesting that other mutations in A.23.1 spike specifically affect fusion in non-TMPRSS2-expressing cells.

The A.23.1 variant pre-dated the B.1.617 lineage as a P681R-containing VOC/VOI by several months. It is interesting to note that B.1.617.2 (Delta) has been shown to be a genetic outlier compared to other VOCs (29), raising the question of whether P681R (found in A.23.1 and B.1.617) ultimately results in a more successful viral variant compared to P681H, found in B.1.1.7 (Alpha) and B.1.529 (Omicron BA.1/BA.2). It is important to note that both lineages that have temporarily dominated Uganda have encoded the spike P681R substitution, but in combination with distinct changes in the spike protein. In all cases, the position 681 change occurred after a change of position 613/614 (B to B.1 to B.1.1.7 (Alpha), B to B.1 to B.1.617.2 (Delta), A to A.23 to A.23.1, B to B.1 to B.1.1.7 to Q.4), and most recently B to B.1 to B.1.1.529 (Omicron). This timing and linkage can be seen in the lineage prevalence charts (Figure 5A and B) where for each major lineage the position 613/614 changes predate the position 681 changes.

The analyses reported here show that the substitution influences furin-mediated cleavage at the *in vitro* level, with these results being consistent with other studies (22, 23). However, P681R may not be the sole driver of spike protein function in vivo—a finding reinforced by the molecular studies described here. It would be of interest to understand the additional spike-associated changes that cooperate with P681R. The introduction of P681R alone into the WT Wuhan-Hu-1 spike did not reproduce the full activity of the A.23.1 spike (Figures 3 and 4). One limitation of this study is that isolated viruses of the A.23.1 lineage are not available for infectivity assays, and so our work relies on the use of epidemiological tools and the reconstruction of virus infection in biochemical and cell-based assays. Another limitation is that our “wild-type” B lineage virus contains D614 and not 614G. Despite these limitations, we consider that our data support our conclusion that the spike mutation P681R—by itself—is a not a primary driver of virus transmissibility in the population, with A.23.1 giving unique insight into these aspects of the ongoing COVID-19 pandemic, but requires the full context of additional spike and other viral changes seen in A.23.1, B.1.617.2 and Q.4 for transmission success.

A.23.1 was a key SARS-CoV-2 variant spreading within Africa during the early part of 2021, and has been defined (along with C.11) as an African VOI (30) having have multiple mutations on the spike glycoprotein and evolving in a clocklike manner along with other variants. Epidemiological data from Uganda support the importance of the P681R substitution in A.23.1 for community-wide transmission. The subsequent decline of the P681R lineage A.23.1 in Uganda, combined with the in vitro analyses reported here clearly showed that the P681R alone is not sufficient to drive such dominance. The P681R lineage B.1.617.2 (Delta) likely benefited from additional S and other substitutions and eventually dominated the Uganda epidemic by June 2021 (Figure 5), similar to patterns globally.

While P681R does make the S1/S2 cleavage site more basic in nature, such variant cleavage sites are still not “ideal” for furin—as originally found in the prototype furin-cleaved virus mouse hepatitis virus (MHV) (RRARR|S) (26, 31). The introduction of an arginine residue did appear to be making S1/S2 more “polybasic” as the pandemic continued and transmissibility increased. While we should not over-simply the complex process of spike protein activation, it will be interesting to see whether this progression of basic residue addition continues with future variants, towards that seen in established community-acquired respiratory coronaviruses such as HCoV-HKU-1 or HCoV-OC43, with S1/S2 sequences of RRKRR|S and RRSRR|A, respectively (26). The recent emergence of B.1.1.529 (Omicron), without P681R but containing distinct changes in its S1/S2 cleavage site (N579K, P681H) and apparently distinct properties in regard to spike protein antigenicity, protease activation and fusion (for example see ref. (32)), has reaffirmed the notion that the coronavirus spike protein is highly adaptable.

## Materials and Methods

### Cells

All cell lines were grown at 37°C with 5% CO_2_. Vero E6 (ATCC CRL-1586) and Hek293T (ATCC CRL-3216) cells were grown in Dulbeco’s Modified Eagle’s Medium (DMEM)(Corning) with 10% HyClone™ FetalClone® II (Cytiva), and 1% HEPES (Corning). Vero TMPRSS2 cells (JCRB Cell Bank JCRB1819) were grown in DMEM, 10% Hyclone, 1% HEPES, and 1% Geneticin (Gibco). A549 ACE2 TMPRSS2 cells (Invivogen a549-hace2tpsa) were maintained in DMEM with 10% FBS, 0.5 μg/ml of Puromycin (Invivogen ant-pr-1) and 300 μg/ml of Hygromycin Gold (Invivogen ant-hg-1).

### Furin prediction calculations

Prop: CoV sequences were analyzed using the ProP 1.0 Server hosted at: https://services.healthtech.dtu.dk/service.php?ProP-1.0. PiTou: CoV sequences were analyzed using the PiTou V3 software hosted at: http://www.nuolan.net/reference.html.

### Fluorogenic peptide assays

Fluorogenic peptide assays were performed as described previously with minor modifications (33). Each reaction was performed in a 100 μl volume consisting of buffer, protease, and SARS-CoV-2 S1/S2 Wuhan-Hu-1 (WT) (TNSPRRARSVA), SARS-CoV-2 S1/S2 B.1.1.7 (TNSHRRARSVA) or SARS-CoV-2 S1/S2 A.23.1 (TNSRRRARSVA) fluorogenic peptide in an opaque 96-well plate. For trypsin catalyzed reactions, 0.8 nM/well TPCK trypsin was diluted in PBS buffer. For furin catalyzed reactions, 1 U/well recombinant furin was diluted in a buffer consisting of 20 mM HEPES, 0.2 mM CaCl2, and 0.2 mM β-mercaptoethanol, at pH 7.5. Fluorescence emission was measured once per minute for 60 continuous minutes using a SpectraMax fluorometer (Molecular Devices) at 30°C with an excitation wavelength of 330 nm and an emission wavelength of 390 nm. Vmax was calculated by fitting the linear rise in fluorescence to the equation of a line.

### Synthesis and cloning of the A.23.1 spike protein

The sequence for the A.23.1 spike gene from isolate SARS-CoV-2 A.23.1 hCoV-19/Uganda/UG185/2020 (EPI_ISL_955136) was obtained from GISAID (https://www.gisaid.org/), codon-optimized, synthesized and cloned into a pcDNA 3.1+ vector for expression (GenScript).

### Site-directed mutagenesis

Primers (ACCTGGCTCTCCTTCGGGAGTTTGTCTGG/CCAGACAAACTCCCGAAGGAGAGCCAGGT) for mutagenesis were designed using the Agilent QuickChange Primer Design tool to create the P681R mutation (CCA->CGA). Mutagenesis was carried out on a pCDNA-SARs2 Wuhan-Hu-1 S plasmid using the Agilent QuickChange Lightning Mutagenesis kit (The original plasmid was generously provided by David Veesler, University of Washington USA). The mutated pcDNA-SARS-CoV-2 Wuhan-Hu-1 P681R S plasmid was used to transform XL-10 gold ultracompetent cells, which were grown up in small culture, and then plasmid was extracted using the Qiagen QIAprep Spin Miniprep Kit. Sanger sequencing was used to confirm the incorporation of the mutation.

### Cell-cell fusion assay

Vero E6 and Vero-TMPRSS2 cells were transfected with a plasmid harboring the spike gene of the SARS-CoV-2 Wuhan-Hu-1 S (WT), SARS-CoV-2 A.23.1 S, SARS-CoV-2 Wuhan-Hu-1 with a P to R mutation in the 681 amino acid position (P681R), or an empty pCDNA3.1+ (S-) plasmid, and evaluated through an immunofluorescence assay (IFA) to quantify nuclei involved in syncytia formation. Transfection was performed on 8-well glass slides at 90% confluent cells using Lipofectamine® 3000 (Cat: L3000075, Invitrogen Co.), following the manufacturer’s instructions and 250 ng of plasmid DNA per well were transfected. Cells were then incubated at 37°C with 5% of CO2 for 28 hours. Syncytia were visualized through fluorescence microscopy using a previously described method (14). The spike expression was detected using a SARS-CoV-2 spike antibody (Cat: 40591-T62, Sino Biological Inc.) at 1/500 dilution for 1 hour. Secondary antibody labeling was performed using AlexaFluorTM 488 goat anti-rabbit IgG antibody (Cat: A32731, Invitrogen Co.) at a 1/500 dilution for 45 minutes. Representative images of each treatment group were used to calculate the percent of nuclei involved in the formation of syncytia. Images were taken at 20X on the Echo Revolve fluorescent microscope (Model: RVL-100-M). Nuclei were counted manually using the Cell Counter plugin in ImageJ (https://imagej.nih.gov/ij/). Cells that expressed the spike protein and contained 4 or more nuclei were considered to be one syncytium.

### Cell surface expression of spike protein

For analysis of cell surface expression Vero E6 and Vero TMPRSS2 cells were seeded at 5×10^5^ cells/ml in a 6 well plate. The following day, each well was transfected using polyethylenimine (PEI) and 1X Gibco® Opti-Mem with 2000 ng of one of the following plasmids: SARS-CoV-2 Wuhan-Hu-1 S (WT), SARS-CoV-2 A.23.1 S, SARS-CoV-2 Wuhan-Hu-1 P681R, or an empty pCDNA3.1+ (S-) plasmid. 24 hours post-transfection, expressed protein on Vero E6 and Vero-TMPRSS2 cells was analyzed through a cell surface biotinylation assay western blot as described previously (34). Spike protein was detected via Western Blot using the antibodies described in the cell-cell fusion assay. GLUT4 protein was used as a housekeeping expression control and labeled using a GLUT4 monoclonal antibody (Cat: MA5-17175, Invitrogen Co.).

### Pseudoparticle generation

Pseudoparticle generation was carried out using a murine leukemia virus (MLV)-based system as previously described with minor modification (18). HEK-293T cells were seeded at 2.5×10^5^ cells/ml in a 6-well plate the day before transfection. Transfection was performed using polyethylenimine (PEI) and 1X Gibco® Opti-Mem (Life Technologies Co.). Cells were transfected with 800ng of pCMV-MLV *gag-pol*, 600ng of pTG-Luc, and 600 ng of a plasmid containing the viral envelope protein of choice. Viral envelope plasmids included pcDNA-SARS-CoV-2 Wuhan-Hu1 S as the WT, pcDNA-SARS-CoV-2 Wuhan-Hu-1 P681R S, and pcDNA-SARS-CoV-2 A.23.1 S. pCAGGS-VSV G was used as a positive control and pCAGGS3.1+ was used for an empty plasmid negative control (S-). 48 hours post-transfection, the supernatant containing the pseudoparticles was removed, centrifuged to remove cell debris, filtered, and stored at -80°C.

### Pseudoparticle Infection Assay

Infection assays were performed as previously described with minor adjustments (18). Vero E6 and Vero-TMPRSS2 cells were seeded at 4.5×10^5^ cells/ml, while A549 ACE2 TMPRSS2 cells were seeded at 4×10^5^ cells/ml in a 24-well plate the day before infection. Cells were washed three times with DPBS and infected with 200 μl of either VSV G, SARS-CoV-2 S, SARS-CoV-2 P681R S, SARS-CoV-2 A.23.1 S, or empty plasmid (S-) pseudoparticles. Infected cells were incubated on a rocker for 1.5 hours at 37°C, then 300 μl of complete media were added and cells were left at 37°C. At 72 hours post-infection, cells were lysed and the level of infection was assessed using the Luciferase Assay System (Cat: E1501, Promega Co.). The manufacturer’s protocol was modified by putting the cells through 3 freeze/thaw cycles after the addition of 100 μl of the lysis reagent. 10 μl of the cell lysate was added to 20 μl of luciferin, and then luciferase activity was measured using the Glomax 20/20 luminometer (Promega Co.). Vero E6 and Vero-TMPRSS2 infection assays were replicated four times. A549 ACE2 TMPRSS2 infection assays were repeated 3 times. Each assay was performed with three technical replicates. Vero E6 and Vero-TMPRSS2 infection assays were completed using the same batch of pseudoparticles, while the A549 ACE2 TMPRSS2 infection assays were carried out using a newly made batch.

### Western blot analysis of pseudoparticles

A 3 ml volume of pseudoparticles was pelleted using a TLA-55 rotor with an Optima-MAX-E ultracentrifuge (Beckman Coulter) for 2 hours at 42,000 rpm at 4°C. untreated particles were resuspended in 30 μl DPBS buffer. Pseudopartices were generated as described in the pseudoparticle generation section, with the furin inhibitor dec-RVKR-CMK (Cat:35-011, Tocris) being added during transfection to select wells. For the + furin treated MLVpps, particles were resuspended in 30 μL of furin buffer consistent in 20 mM HEPES, 0.2 mM CaCl2, and 0.2 mM β-mercaptoethanol (at pH 7.0). Furin-treated particles were later incubated with 6 U of recombinant furin for 3 h at 37 °C. Sodium dodecyl sulfate (SDS) loading buffer and DTT were added to all samples and heated at 95°C for 10 minutes. Samples were separated on NuPAGE Bis-Tris gel (Invitrogen) and transferred on polyvinylidene difluoride membranes (GE). SARS-CoV-2 S was detected using a rabbit polyclonal antibody against the S2 domain (Cat: 40590-T62, Sinobiological) and an AlexaFluor 488 goat anti-rabbit antibody. Bands were detected using the ChemiDoc Imaging software (Bio-Rad) and band intensity was calculated using the analysis tools on Biorad Image Lab 6.1 software to determine the uncleaved to cleaved S ratios.

### Uganda cases vs. viral lineages over time

Daily reported SARS-CoV-2 infections The Uganda daily SARS-CoV-2 positive samples numbers were retrieved from Our World in Data (https://ourworldindata.org/coronavirus) and the 7 day average was determined. Uganda SARS-CoV-2 lineage data were generated from the MRC Uganda genomic data deposited in the GISAID database (https://www.gisaid.org/). SARS-CoV-2 Pango lineages (2) were determined using the pangolin module pangoLEARN (https://github.com/cov-lineages/pangolin).

### Spike position 681 and 613/614 changes in global data

All available spike protein sequences were obtained from the GISAID database. The frequency of P681, P681R, P681H, D614 D614G and D613H were counted by string matching using Ack (http://beyondgrep.com/) to the major variations of the 88 amino acid peptide sequence (aa 605 to 691) spanning the two relevant sites (D614_P681 (Wuhan_B), D614G_P681 (B.1), D614_P681H, D614G_P681H (B.1.1.7), D614_P681R, Q613_P681 (Wuhan_B), Q613H_P681 (A.23), Q613H_P681R (A.23.1), D614G_P681R (B.1.617.2). Fractions of available total genomes for each month encoding each peptide variant were visualized in a heatmap. Additional changes at position H655Y (present in the Gamma lineage) were also included in the count and fraction calculation but the total numbers were minor.

### Quantification and Statistical Analysis

All statistical analysis was performed using GraphPad Prism for Mac OS X, GraphPad Software, San Diego, California USA, www.graphpad.com. Two sample T-tests were used to compare SARS-CoV-2 Wuhan-Hu 1 to SARS-CoV-2 A.23.1 or SARS-CoV-2 P681R, with significant P values reported in the figure legends. Standard deviation was calculated and included in graphs when appropriate.

## Acknowledgements

This work was funded in part by the National Institute of Health research grant R01AI35270 (to GW and SD). We thank the global SARS-CoV-2 sequencing groups for their open and rapid sharing of sequence data and GISAID for providing an effective platform to make these data available. DLB, MVTP and MC were funded by the UK Medical Research Council (MRC/UK Research and Innovation) and the UK Department for International Development (DFID) under the MRC/DFID Concordat agreement (grant agreement no. NC_PC_19060) and Wellcome Trust, UK FCDO—Wellcome Epidemic Preparedness—Coronavirus (grant agreement no. 220977/Z/20/Z). TT was supported by the National Science Foundation Graduate Research Fellowship Program under Grant No. DGE-1650441 and the Samuel C. Fleming Family Graduate Fellowship.

## Author contributions

Conceptualization: BL, LEF, TT, SD, JAJ and GRW. Methodology: BL, LEF, MVTP, DLB, TT, SD, MC, JAJ and GRW. Investigation: BL, LEF, MVTP, DLB, TT, JAJ. Writing - Original Draft: BL, TT, JAJ. and GRW. Writing - Review & Editing, BL, MVTP, TT, SD, MC, JAJ and GRW. Visualization: BL, LEF, MVTP, TT and JAJ. Supervision: SD, MC, JAJ and G.R.W. Funding acquisition: SD, MC and GRW.

